# libSBOLj3: A graph-based library for design and data exchange in synthetic biology

**DOI:** 10.1101/2023.05.19.540930

**Authors:** Göksel Mısırlı

## Abstract

The Synthetic Biology Open Language version 3 data standard provides a graph-based approach to exchange information about biological designs. The new data model has major updates and offers several features for software tools. Here, we present libSBOLj3 to facilitate data exchange and provide interoperability between computer-aided design and automation tools using this standard. The library adopts a graph-based approach. Tool developers can extend these graphs with application-specific information and use detailed validation reports to identify errors and interoperability issues and apply best practice rules. The libSBOLj3 library is implemented in Java and can be downloaded or used as a Maven dependency. The open-source project, examples and documentation about accessing and using the library are available via GitHub at https://github.com/SynBioDex/libSBOLj3.

## Introduction

The standard representation of data is ever more important for reproducibility and the development of predictable applications for synthetic biology. Data standards facilitate taking advantage of different computer-aided design and automation tools, interoperability between these tools and the development of complex workflows^1^.

The Synthetic Biology Open Language (SBOL) is an open-source data standard to facilitate the electronic exchange of information about genetic parts, designs and related information. Several genetic design, visualisation, modelling and automation tools use SBOL to represent and exchange data^2^. Moreover, data repositories adopt SBOL for data storage, integration and retrieval^3^.

The initial version of SBOL focused on representing structural information at the DNA level for modular and hierarchical reuse of genetic parts. SBOL’s data model was then extended for different types of parts such as RNA, protein and small molecules and molecular interactions, facilitating the reuse of designs with well-defined inputs and outputs^4^.

The recently developed SBOL3^5^ adopts a fully graph-based approach in which SBOL entities represent nodes, and edges represent relationships between these entities. A linked data approach is used to identify and describe SBOL entities via existing ontologies and controlled vocabularies^6–8^. SBOL3 has also been extended to support synthetic biology workflows more efficiently to build a web of design information.

Here, we present libSBOLj3, a Java library for SBOL3. The library is document-centric and uses a graph-based approach to handle SBOL graphs efficiently. Tool developers can use this library to take advantage of the recent SBOL3 data standard, which provides increased reuse of biological design information and richer expressions to represent design-related constraints. Tool developers can also incorporate application-specific information and describe the provenance-based design, build and test activities.

## Results

The libSBOLj3 provides several features. Some of the important features are summarised below.

### Document-centric data exchange

Each SBOL document can have multiple entities designed to represent information of interest, such as a `Component` entity to represent a genetic part and a `Sequence` entity to define the composition of a part. These top-level container entities are further described using child entities (Figure 1). The library can write the resulting SBOL documents into files or in-memory variables and read them back.

**Figure 1:**
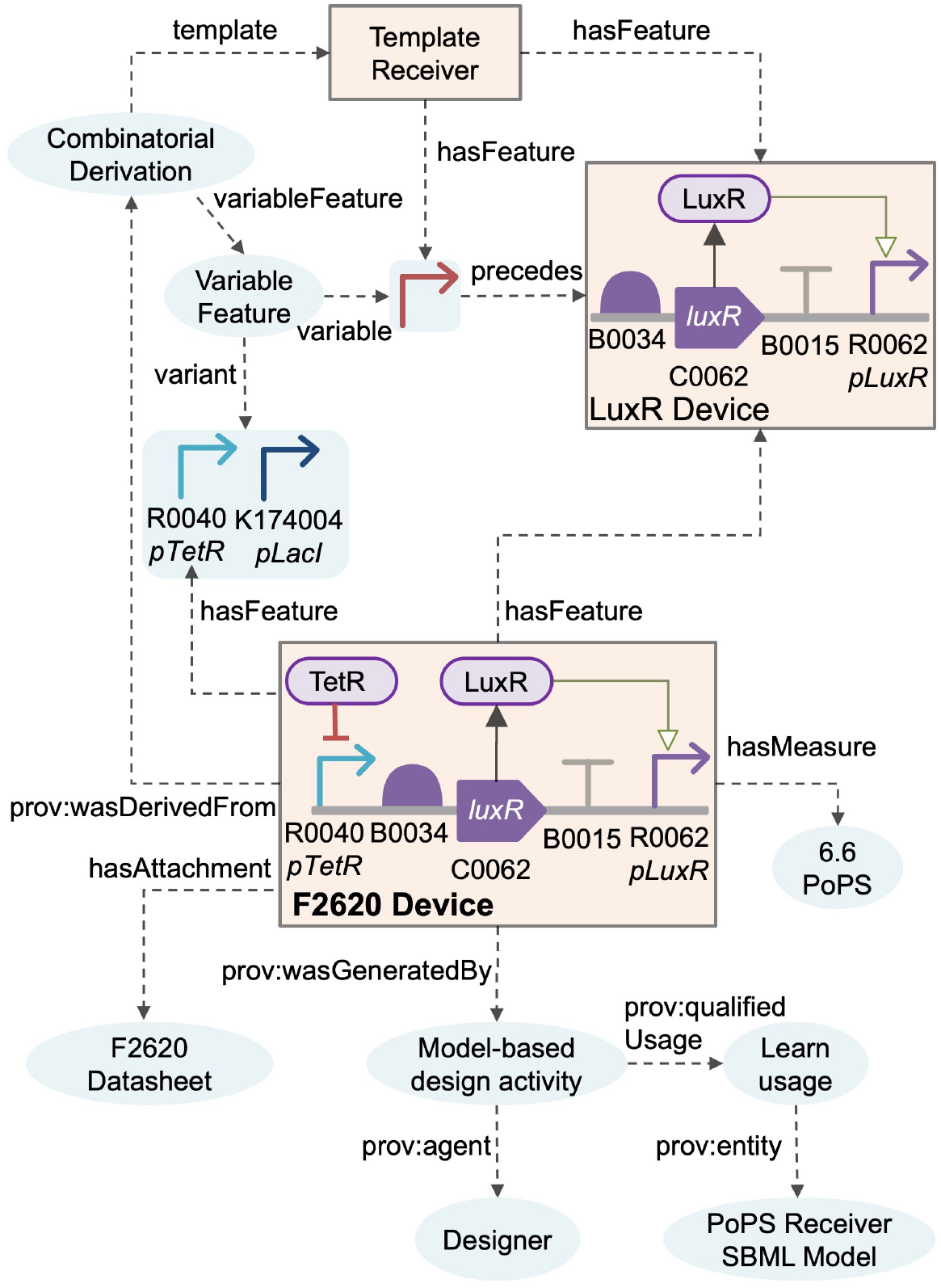
An example libSBOLj3 document that represents a genetic circuit design and related information as a biological network. The F2020 design is derived from a template with well-defined features that can be varied combinatorially. Functional constraints about information about molecular interactions and structural constraints such as precedes and overlaps are incorporated. A datasheet about experimental results and provenance information about how this device is derived using a model-based design activity are associated with the design. A measurement entity with a polymerases per second (PoPS) value is also included.

### Graph-based documents

The library associates each SBOL document with a graph and hides the complexity of working with these graphs. Different serialisation formats can be used when writing or reading SBOL graphs, which can also be processed by off-the-shelf graph tools to store, visualise and query data.

- `RDF/XML` to support graph and XML tools.
- `Turtle` as a human-readable format.
- `JSON-LD` to use lightweight JSON graphs for web applications.
- `N-Triples`, when ordered by libSBOLj3, can facilitate versioning, comparing and streaming SBOL documents.

### Object-oriented application programming interface

The libSBOLj3 library adopts an object-oriented data manipulation and retrieval mechanism. Programmatic retrieval and update of SBOL entities and their properties are via simple getters and setters. Each SBOL object includes a reference to a graph node, and object properties and child objects are accessed when needed to support large and complex SBOL graphs. This lazy loading approach minimises the unnecessary interaction with the underlying graph and the creation of objects until these objects are requested. When an SBOL property is updated, the corresponding graph node is also updated.

The library provides additional graph-based access mechanisms for SBOL entities. An entity can be retrieved using its unique graph identifier via the `getIdentified` library call. The library also supports graph queries to search for entities. The search language is built on SPARQL SELECT. Prefix definitions and the SELECT clause are omitted, and a single search variable corresponding to SBOL entities with given criteria is used. For example, the `getIdentifieds` method returns terminator genetic parts when it is executed with the “`?identified a sbol:Component; sbol:role SO:0000141; sbol:type SBO:0000251 .`” query as input, and a list of Component objects is returned.

The library provides methods to control the creation of child entities with valid identifiers according to SBOL rules while offering flexibility to improve developers’ user experience. Example SBOL rules include that child entities’ identifiers must start with their parents’ identifiers, and top-level entities’ identifiers must not be included in other top-level entities. An SBOL document provides factory methods to create top-level entities using the document’s base identifier consistently. Similarly, an SBOL entity provides methods to create valid child entities.

### High-level SBOL API

The library provides a high-level application programming interface (API) for repetitive or complex tasks and to decouple implementation details from the SBOL data model. Such methods can be used when design information relies on various entities and external ontologies or when describing complex workflows. These higher-level methods are regularly added based on community feedback.

### Application-specific data support

The libSBOLj3 library supports integrating application-specific data using a linked data approach. Custom data can be included as top-level or child entities or used to annotate existing SBOL entities in the form of properties and values. The values corresponding to non-SBOL properties can be literals or identifiers for external or internal entities.

### Validation

Validation is an essential feature of the library to create valid SBOL documents. SBOL documents are validated for required and conformance-related rules and best practices. Validating best practices can be switched off programmatically. The library can validate an SBOL document, an SBOL entity or a folder of SBOL documents. Only valid documents can be serialised. Invalid values and entities are reported with a list of validation messages which rely on SBOL-specific validation codes and descriptions. Contextual information about how an invalid value relates to a validated entity may involve a subgraph when multiple entities are involved. Such a subgraph is defined textually as a path between the validated entity and the invalid value in the form of a set of property names and values.

## Methods

The libSBOLj3 library has been developed using the Java programming language. The library builds upon Resource Description Framework (RDF) graphs^9^. The graph abstraction is achieved via the Jena^10^ library, and the validation layer is built on Hibernate Validator^11^. Project-specific dependencies and build automation are controlled with Maven ^12^. Test-driven development was adopted to create a user-friendly library to test validation rules effectively, providing more than 95% code coverage in unit tests.

## Conclusion

The library presented here has been developed to facilitate interoperability between software tools in synthetic biology. Whilst the library can be used to work directly with the SBOL data model, the high-level API provides intuitive and less verbose methods to increase the user experience. Tool developers can use this library to create valid SBOL documents without the complexity of dealing with graphs and can take advantage of graph-based biological data integration, processing and querying.

## Acknowledgements

We thank the SBOL community, SBOL3 Working Group, Jacob Beal, Tom Mitchell and James McLaughlin for their valuable comments and Chris Myers for testing the library and providing feedback regularly. We thank Keele University for Research Development Fund and James Rooney, Andrew Cook and Rory Gee for their help in testing and documentation.

## Data availability

The data underlying this article are available on GitHub at https://github.com/SynBioDex/libSBOLj3.

